# A human-ACE2 knock-in mouse model for SARS-CoV-2 infection recapitulates respiratory disorders but avoids neurological disease associated with the transgenic K18-hACE2 model

**DOI:** 10.1101/2024.06.11.598471

**Authors:** Anna Pons-Grífols, Ferran Tarrés-Freixas, Mònica Pérez, Eva Riveira-Muñoz, Dàlia Raïch-Regué, Daniel Pérez-Zsolt, Jordana Muñoz-Basagoiti, Barbara Tondelli, Edwards Pradenas, Nuria Izquierdo-Useros, Sara Capdevila, Júlia Vergara-Alert, Victor Urrea, Jorge Carrillo, Ester Ballana, Stephen Forrow, Bonaventura Clotet, Joaquim Segalés, Benjamin Trinité, Julià Blanco

**Author notes:** Corresponding authors Benjamin Trinité Julià Blanco. JS, BT and JB are co-senior authors.

## Abstract

Animal models have been instrumental in elucidating the pathogenesis of SARS-CoV-2 infection and testing COVID-19 vaccines and therapeutics. Wild-type (WT) mice are not susceptible to many SARS-CoV-2 variants, therefore transgenic K18-hACE2 mice have emerged as a standard model system. However, this model is characterized by severe disease, particularly associated with neuroinfection, which leads to early humane endpoint euthanasia. Here, we established a novel knock-in (KI) mouse model by inserting the original K18-hACE2 transgene into the collagen COL1A1 locus using a recombinase mediated cassette exchange (RMCE) system. Once the Col1a1-K18-hACE2 mouse colony was established, animals were challenged with a B.1 SARS-CoV-2 (D614G) isolate and were monitored for up to 14 days. Col1a1-K18-hACE2 mice exhibited an initial weight loss similar to the K18-hACE2 transgenic model but did not develop evident neurologic clinical signs. The majority of Col1a1-K18-hACE2 mice did not reach the preestablished humane endpoint, showing progressive weight gain after 9 days post-infection (dpi). Importantly, despite this apparent milder pathogenicity of the virus in this mouse model compared to the K18-hACE2 transgenic model, high levels of viral RNA were detected in lungs, oropharyngeal swab, and nasal turbinate. Remaining lesions and inflammation in lungs were still observed after 14 dpi. In contrast, although low level viral RNA could be detected in a minority of Col1a1-K18- hACE2 animals, no brain lesions were observed at any timepoint. Overall, Col1a1-K18- hACE2 mice constitute a new model for investigating SARS-CoV-2 pathogenesis and treatments, with potential implications for studying long-term COVID-19 sequelae.

**Importance:** K18-hACE2 mice express high levels of the human protein ACE2, the receptor for SARS-CoV-2, and can therefore be infected by this virus. These animals have been crucial to understand viral pathogenesis and to test COVID-19 vaccines and antiviral drugs. However, K18-hACE2 often die after infection with initial SARS-CoV-2 variants likely due to a massive brain infection that does not occur in humans. Here, we used a technology known as knock-in that allows for the targeted insertion of a gene into a mouse and we have generated a new hACE2-mouse. We have characterized this new animal model demonstrating that the virus replicates in the respiratory tract, damaging lung tissue and causing inflammation. In contrast to K18-hACE2 mice, only limited or no brain infection could be detected, and most animals recovered from infection with remaining lung lesions. This new model could be instrumental for the study of specific disease aspects such as post-COVID condition, sequelae, and susceptibility to reinfection.

## Introduction

Severe Acute Respiratory Syndrome Coronavirus 2 (SARS-CoV-2) is the etiologic agent of the coronavirus disease 2019 (COVID-19). The COVID-19 pandemic fueled an unprecedented collaborative research effort to develop therapeutics and vaccines. Animal models that recapitulate key clinical and pathological features of COVID-19 have played a pivotal role in testing novel vaccines, antivirals and other treatments (1).

The spike glycoprotein of SARS-CoV-2 uses the angiotensin-converting enzyme 2 (ACE2) as receptor to enter and infect host cells (2). ACE2 is expressed on the surface of cells of different organs such as lungs, intestines, kidneys, and heart. However, the spike glycoprotein from ancestral SARS-CoV-2 strains does not efficiently bind to mouse ACE2 (mACE2), rendering wild-type (WT) mice refractory to infection. Therefore, several strategies were used to develop mouse models susceptible to infection. These strategies include viral adaptation (3) or the introduction of the human ACE2 (hACE2) receptor via viral vectors and transgenic approaches (4–8).

Both K18-hACE2 transgenic mice and Golden Syrian hamsters (GSH) became reference animal models for investigating SARS-CoV-2 pathogenesis in vivo. They mainly differ in pathogenesis: GSH, which are naturally susceptible to SARS-CoV2 infection, recapitulate a milder disease phenotype (1). The K18-hACE2 mouse model was originally developed for the study of SARS-CoV, which also targets hACE2. It was created by random insertion of multiple copies of the *hACE2* gene under the control of the human cytokeratin 18 gene promoter (*KRT18* or K18). This promoter allows for high-level expression and is specific to epithelial cells, including those in the airways (9, 10). This approach allows the infection of mice by SARS-CoV-2, while maintaining the activity of the mACE2 receptor. Infection of K18- hACE2 mice with pre-Omicron variants of SARS-CoV-2 results in progressive weight-loss and strong clinical signs by 3-5 days post-infection (dpi), a severity which often requires euthanasia by 5-7 dpi. The lethality in this model is dose- and SARS-CoV-2 variant-dependent (10, 11), and has been associated with neuroinvasion and extensive brain infection (9, 12–14). Neuroinvasion in K18-hACE2 transgenic mice has been linked to the aleatory gene insertion and the high number of inserted gene copies, around 8 in the commercial *B6.Cg-Tg(K18-ACE2)2Prlmn/J-K18-hACE2 mouse* (10), leading to altered expression levels and tissue distribution of the receptor (15). Specifically, a higher number of gene copies correlates with a worse disease prognosis and encephalitis in this model, following SARS-CoV infection (16). These neurological lesions leading to death are not consistent with the pathology of COVID-19 in humans, in which infection of the central nervous system can be detected post-mortem in severe COVID-19 cases, although to a lower extent and unrelated to the severity of brain lesions (17, 18). Therefore, new mouse models that better recapitulate the human disease, without lethal viral neuroinvasion, could complement other existing animal models (hamster or NHP) (1).

Knock-in (KI) models are characterized by the targeted insertion of a defined gene copy number in a specific locus, therefore allowing for a more accurate and predictable expression of the transgene. In this work, we used a recombinase-based approach to insert the original K18-hACE2 transgene into the collagen type I alpha chain (COL1A1) locus to generate a Col1a1-K18-hACE2 KI mouse. To characterize this new model, mice were challenged with SARS-CoV-2 B.1 (D614G) isolate and pathogenicity was compared with the well-characterized K18-hACE2 transgenic mice. After viral challenge of Col1a1-K18-hACE2 mice, we confirmed viral replication and histological lesions in the lungs, but minimal viral neuroinvasion.

## Results

### Assessment of hACE2 expression in the Col1a1-K18-hACE2 mice

We generated a new hACE2 KI mouse model through addition of the original K18-hACE2 transgene, kindly supplied by Paul McCray (9) at the COL1A1 locus using a recombinase mediated system (**Figure 1A**) (19). Traditional transgenic techniques rely on the random genomic insertion of an unpredictable number of transgene copies. This frequently results in a highly variable level and pattern of transgene expression, due to cis acting regulatory elements at the site of insertion. A single copy of a transgene, inserted into the well characterised Col1A1 locus, is expected to result in a more predictable level and pattern of expression of the transgene sequence. To confirm the correct expression of the transgene in this KI model, we initially quantified *hACE2* transcripts relative to the house-keeping gene GAPDH by RT-qPCR in RNA samples extracted from twelve different tissues. Expression of the hACE2 mRNA in K18-hACE2 mice was similar among most tested tissues except in muscle and heart, which showed lower values (**Figure 1B).** In contrast, hACE2 mRNA expression in Col1a1-K18-hACE2 mice was more heterogenous, with the lowest expression observed in the brain and liver. When comparing the relative expression of hACE2 mRNA between both models, we observed higher expression in the pancreas, lymph nodes and muscle of Col1a1-K18-hACE2 mice. Conversely, in the kidney, brain and liver, hACE2 mRNA expression was higher in the K18-hACE2 mouse model. Minor differences were observed in the rest of the analyzed tissues (**Figure 1C**). To complement mRNA expression data, protein expression was assessed by Wester-blot (WB) in 6 different tissues using a monoclonal antibody specific for hACE2 (20) and a polyclonal antibody recognizing both hACE2 and mACE2. GAPDH was used as control, and tissues from Col1a1-K18-hACE2, K18-hACE2 and WT mice were analyzed. Data showed undetectable protein expression in muscle, and higher hACE2 expression in lung, pancreas and spleen of Col1a1-K18-hACE2 mice compared to K18-hACE2 animals **(Figure 1D)**. Conversely, as observed by RT-qPCR, a higher hACE2 expression was detected in K18-hACE2 brain and liver compared to Col1a1-K18-hACE2 samples **(Figure 1D)**.

**Figure 1.**
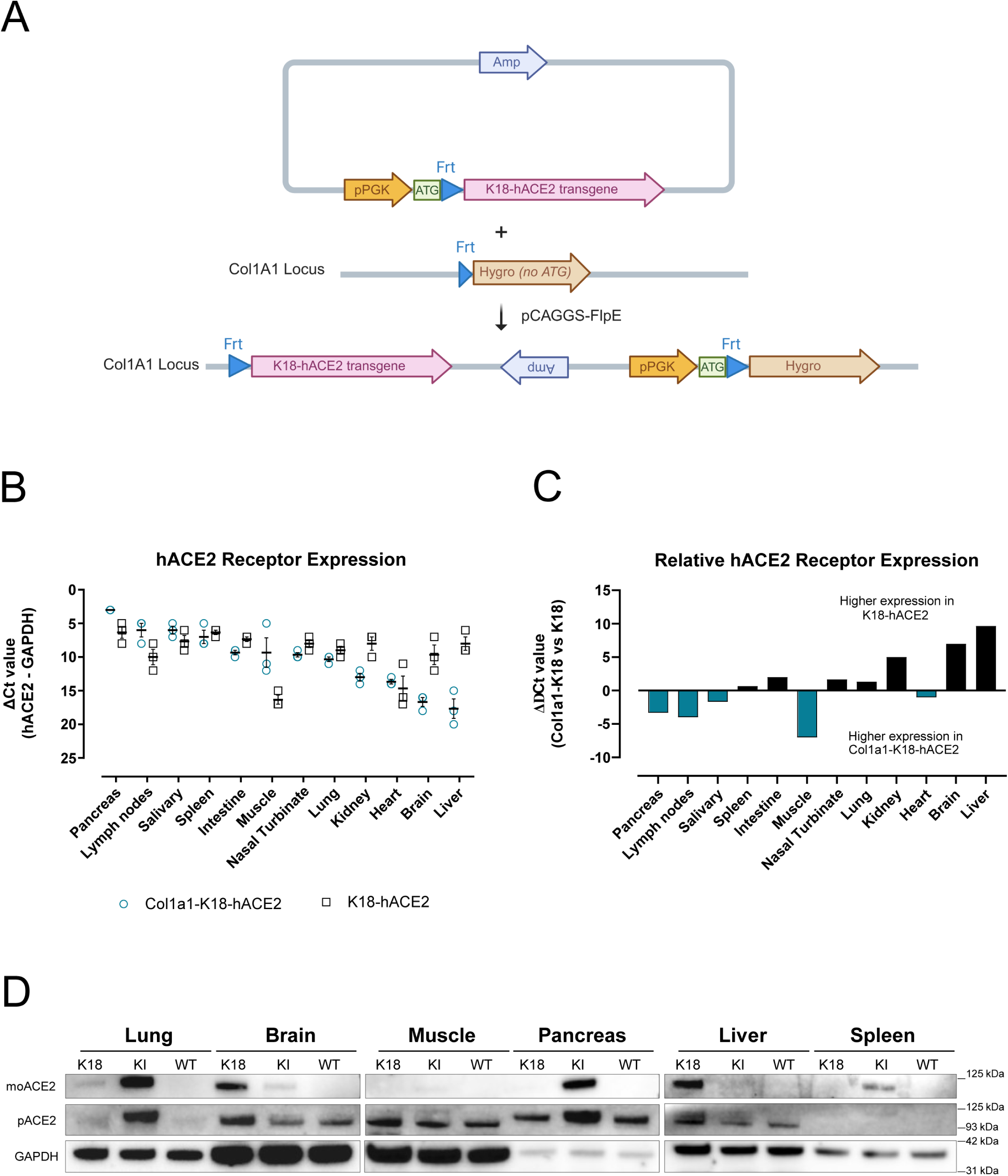
Expression of hACE2 in the Col1a1-K18-hACE2 KI model. (A) Schematic representation of the insertion strategy. The original K18-hACE2 transgene was inserted into the collagen COL1A1 locus using a recombinase mediated cassette exchange (RMCE) FLP-FRT system in KH2 cells via blastocyst injection. pPGK-ATG-frt plasmid: vector backbone with Ampicillin resistant gene (Amp), transcription start site (ATG) and Flippase recognition target (Frt). Hygro: Hygromicin resistance gene. pCAGGS-FlpE: expression plasmid for FLPe recombinase expression. (B) Relative quantification of hACE2 receptor expression to GAPDH expression in uninfected Col1a1-K18-hACE2 ((blue empty dot, n=3) and K18-hACE2 males (black empty square, n=3). Delta Ct values are inversely shown to facilitate interpretation. The lower the absolute number, the higher the relative expression. Solid line and bars represent mean and SEM. (C) Relative comparison of hACE2 receptor expression in Col1a1- K18-hACE2 versus K18-hACE2 mice (n=3 each, males). Higher receptor expression in Col1a1- K18-hACE2 (negative values in Y axis) is marked by blue bars, and higher expression in K18- hACE2 (positive values in Y axis) is marked in black bars. (D) Western Blot analysis of hACE2, mACE2 and GAPDH in representative tissues. hACE2 signal was obtained using a specific monoclonal antibody (top panels); mACE2 and hACE2 signal was obtained with a cross-reactive polyclonal antibody (middle panel), and anti-GAPDH was analyzed as reference (bottom panels). Molecular weight marker is shown on the right.

### Survival of Col1a1-K18-hACE2 mice following SARS-CoV-2 B.1 infection

Col1a1-K18-hACE2 mice (n=27) and K18-hACE2 mice (n=4) were challenged intranasally with a SARS-CoV-2 B.1 (D614G) isolate. Uninfected KI animals, used as a control, received an intranasal dose of PBS (n=5). Animal weight and clinical signs were monitored daily for 14 days after infection. The endpoints to collect samples were set at 3-, 7-, and 14-days post-infection (dpi, n=8 Col1a1-K18-hACE2 mice per endpoint) or upon fulfilment of humane endpoint criteria (**Figure 2A**).

**Figure 2.**
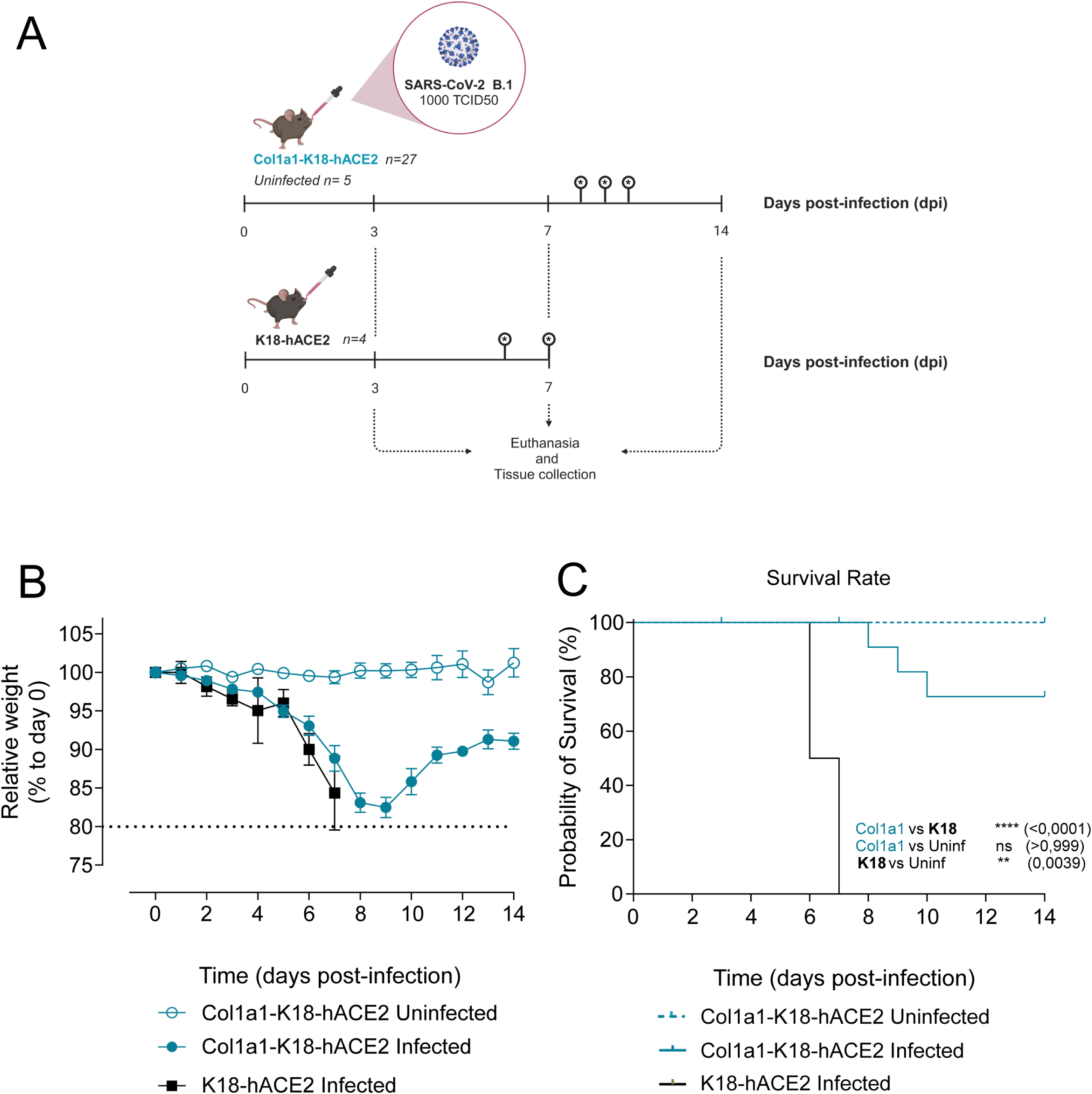
Experimental setting and progression of SARS-CoV-2 infection in Col1a1-K18- hACE2 and K18-hACE2 mouse models. (A) Schematic representation of the experimental setting. Knock-in Col1a1-K18-hACE2 (n=27) and transgenic K18-hACE2 mice (n=4) were intranasally challenged with a 1000 TCID_50_ dose of a B.1 SARS-CoV-2 isolate. A Col1a1-K18- hACE2 uninfected control group (n=5) was challenged with PBS. Mice were monitored for weight-loss and clinical signs for 14 days post infection (dpi). Euthanasia was performed at 3, 7, and 14 dpi or upon fulfilment of humane endpoint criteria, for sample and tissue collection (n=8 per timepoint). Infections were performed in two separate experiments between January and June 2022. Created with Biorender.com. (B) Relative body weight follow-up referred to day 0. Col1a1-K18-hACE2 uninfected (blue empty dot), Col1a1-K18- hACE2 infected (blue dot), K18-hACE2 infected (black square). Solid lines and bars represent mean±SD. (C) Survival (Kaplan-Meier). All K18-hACE2 infected animals (n=4) had to be euthanized due to endpoint criteria by 7dpi, and only three infected Col1a1-K18-hACE2 at 8, 9, and 10 dpi. No uninfected Col1a1-K18-hACE2 had to be euthanized under this criterion. Col1a1-K18-hACE2 uninfected (blue dashed line), Col1a1-K18-hACE2 infected (blue line), K18-hACE2 infected (black line). Statistical differences were identified using a Log-rank (Mantel-Cox) test (<0.0001), followed by individual comparisons (**p < 0.005; ****p < 0.0001)

Consistent with previous studies in transgenic animals with the B.1 SARS-CoV-2 variant, weight-loss could be observed in both animal models starting around 3 dpi (**Figure 2B**) (13). In K18-hACE2 transgenic animals, the decrease in weight was associated with the progressive appearance of severe clinical signs: decrease in mobility, dyspnea, and neurological effects **(Supplementary Table 1),** fulfilling the humane endpoint criteria and leading to euthanasia of all K18-hACE animals by 6 or 7dpi (**Figure 2C**). In contrast, while Col1a1-K18-hACE2 mice underwent weight loss, they did not show severe clinical signs apart from a slight decrease in activity. Three Col1a1-K18-hACE2 animals had to be euthanized (exitus) exclusively due to their weight loss exceeding the 20% limit (humane endpoint) at 8, 9 and 10 dpi. Convalescent animals started to regain weight after 9 dpi, reaching 90% of their initial weight by 14 dpi (**Figure 2B/C**). These results indicate that B.1 SARS-CoV-2 infection is significantly less pathogenic in Col1a1-K18-hACE2 mice than in K18- hACE2 transgenic mice (**Figure 2C**).

### SARS-CoV-2 B.1 replicates in the respiratory tract of Col1a1-K18-hACE2 KI mice

To fully characterize the kinetics of viral replication in the respiratory tract, samples from oropharyngeal swab, lung, and nasal turbinate were collected at 3, 7, and 14 dpi or at the humane endpoint (exitus) in all Col1a1-K18-hACE2 mice (n=32). Samples from K18-hACE2 transgenic mice obtained at euthanasia (6 and 7 dpi) were used as reference (n=4). Viral RNA analysis in oropharyngeal swab, lung, and nasal turbinate revealed widespread infection in both K18-hACE2 and Col1a1-K18-hACE2 mice in all mentioned tissues across two independent experiments (**Figure 3A**). VL in the nasal turbinate were similar in both animal models, with some viral RNA was still detected at 14 dpi in Col1a1-K18-hACE2 mice. RNA levels in oropharyngeal swabs and lung peaked at 3 dpi in Col1a1-K18-hACE2 mice and tended to decay overtime up to 14 dpi. To further confirm viral clearance, titration of replication-competent virus was performed in lung samples. Similar to the VL kinetics, viral titers peaked at 3 dpi, followed by a quick and significant decrease (**Figure 3B**, 3 dpi vs 7 dpi: *p* = 0.00178), concomitant with the elicitation of neutralizing antibodies **(Supplementary** Figure 1). Col1a1-K18-hACE2 animals euthanized at the humane endpoint at 8-10 dpi still displayed high viral RNA in lungs, which could explain their more pronounced weight-loss and early fulfilment of endpoint criteria. However, high titers of neutralizing antibodies in these animals probably prevented the recovery of infectious viruses **(Figure 3B and Supplementary** Fig 1).

**Figure 3.**
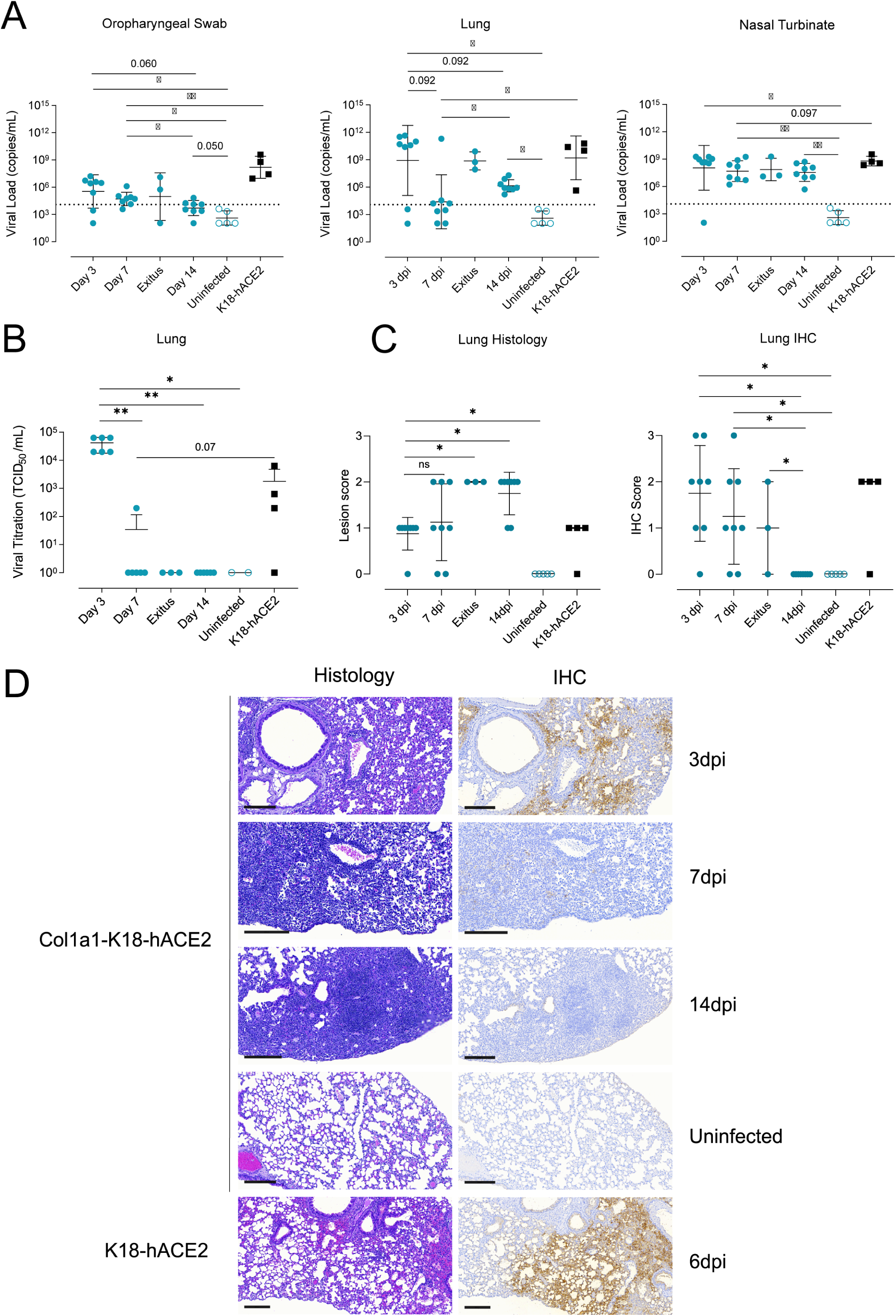
Progression of SARS-CoV-2 infection in Col1a1-K18-hACE2 KI mice. Col1a1-K18- hACE2 (blue circles, *n=27*) and K18-hACE2+ mice (black squares, *n=4*) were inoculated with 1000 TCID_50_ of a SARS-CoV-2 B.1 isolate (full shapes) or uninfected (empty shapes of each colour, *n=5*). Animals were euthanized at 3dpi (n=8), 7dpi (n=8), 14 dpi (n=8) or upon fulfilment of humane endpoint criteria **(n=3, euthanized at 8,9 and 10 dpi)**. (A) SARS-CoV-2 viral RNA loads (copies/mL) of oropharyngeal swab, lung, and nasal turbinate samples. Dashed line represents limit of detection, established by 2SD of uninfected animals. Statistical differences were identified using a Peto-Peto Left censored test (*p < 0.05, **p<0.005). (B) Viral titration of replicative virus (TCID_50/_mL) in lung samples from B.1 infected mice at different endpoints in Vero E6 cells on day 5 of culture. Titers were compared using an Independence Asymptotic Generalized Pearson Chi-Squared Test for ordinal data (*p < 0.05, **p<0.005). (C) Lung histopathological scoring of broncho- interstitial pneumonia (left) and SARS-CoV-2 NP immunohistochemical (IHC) scoring (right) in both models at 3, 7, 14 dpi and endpoint. Statistical differences were identified using an Independence Asymptotic Generalized Pearson Chi-Squared Test for ordinal data (*p < 0.05). (D) Representative lung histology and IHC pictures of both models at 3, 7, 14 dpi and endpoint. Images show low-power magnification bars (200 µm).

To characterize histopathological changes, formalin-fixed lung samples from both animal models were analyzed by histology and immunohistochemistry (IHC) to score the presence and extension of lesions and SARS-CoV-2 nucleoprotein antigen (NP), respectively. Histological lesions were semi-quantified as none, mild, moderate, or severe (score 0-3), and the detection of viral antigen as lack, low, moderate, or high (score 0-3). In Col1a1-K18- hACE2 KI mice, histopathological analysis showed development of bronchointerstitial pneumonia characterized by multifocal increased thickness of interalveolar wall, presence of macrophage-like cells into alveoli surrounding bronchi and bronchiole, and hyperplasia of type II pneumocytes. These lesions evolved from mild and multifocal by 3 dpi to moderate in most of the animals by 14 dpi, also with increasing lymphoplasmacytic infiltration until the end of the study. The extension of hyperplasia of type II pneumocytes was barely detected at 3 dpi, mild by 7 dpi and moderate by 14 dpi. Therefore, an increase in lung lesion average was observed over time **(Figure 3C and D).** Animals reaching humane endpoint displayed areas of severe bronchointerstitial pneumonia (**Supplementary** Figure 2). IHC detection of SARS-CoV-2 NP in the lungs of Col1a1-K18-hACE2 KI mice was consistent with the VL analyses, showing the highest score early upon infection (3 dpi). Antigen detection decreased with time showing apparent clearance by 14 dpi **(Figure 3C and D).** However, animals reaching humane endpoint at 8,9 and 10 dpi showed mild to moderate SARS-CoV- 2 antigen staining mainly located in bronchiolar epithelium from different bronchiole (**Supplementary** Figure 2).

To assess infection-driven inflammation, we analyzed the levels of IL-6 and IFNγ cytokines, involved in inflammation and immune activation; and IP-10, MCP-1 and MIP-1β chemokines, primarily involved in the recruitment of immune cells. Cytokine and chemokine levels were measured by Luminex in lung samples from both animal models at the indicated timepoints. While all markers were increased in infected animals, compared to uninfected controls, IP-10 and IL-6 peaked at 3 dpi and significantly decreased by 14 dpi in Col1a1-K18- hACE2 KI mice (**Figure 4**). In contrast, IFNγ, MCP-1, and MIP1β peaked at 7 dpi and decayed afterwards, although no statistical differences were observed among the analyzed timepoints. Overall, levels of inflammatory mediators at 7 dpi in Col1a1-K18-hACE2 KI mice were comparable to K18-hACE2 transgenic mice at the time of euthanasia (6 and 7 dpi). Residual levels of IFNγ and MIP1β were observed at 14dpi in some Col1a1-K18-hACE2 mice, probably associated with the remaining tissue lesions at this time point, since no viral antigen could be detected **(Figure 4. C-D).**

**Figure 4.**
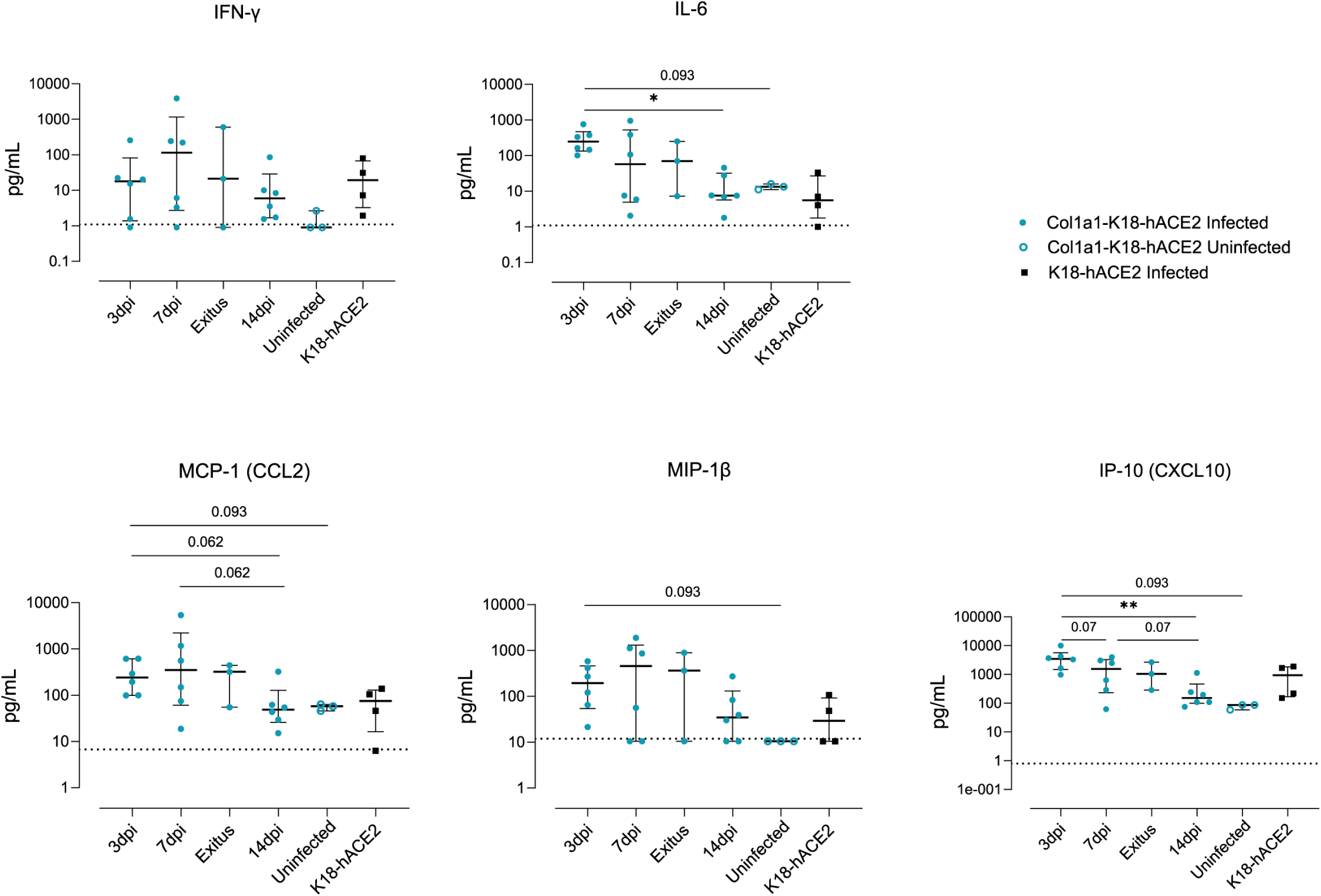
Inflammatory response in the lung. The concentration of Inflammatory cytokines in lung extracts in the knock-in and transgenic models at 3,7,14 dpi or endpoint are shown. Col1a1-K18-hACE2 infected (blue full circles, *n=23*), uninfected (blue empty circles, n=3) and K18-hACE2 infected mice (black full squares, *n=4*). The bar shows Median with interquartile range. The limit of detection for each cytokine is indicated by a dotted line. Statistical differences were identified using a Kruskal Wallis, and a Conover’s nonparametric all-pairs comparison test (*p < 0.05, **p<0.005).

### SARS-CoV-2 B.1 replication in other tissues

To explore the extent of viral infection in both animal models, viral loads (VL, cell-free viral RNA) were quantified in a large set of samples (brain, muscle, intestine, liver, kidney, pancreas, salivary glands, lymph nodes and spleen) collected from a subset of animals (n=20). In both models, VL were mostly undetected in most tissues except for low levels found in the heart and salivary glands (**Figure 5**). The analysis of brain samples showed high VL exclusively in K18-hACE2 transgenic animals. However, the initial screening of brain tissue from Col1a1-K18-hACE2 mice showed low but detectable VL in one animal at 3 dpi and two animals at 7 dpi. In sharp contrast, euthanized animals, which showed the highest weight loss and lung VL, lacked detectable VL in the brain (**Figure 5**).

**Figure 5.**
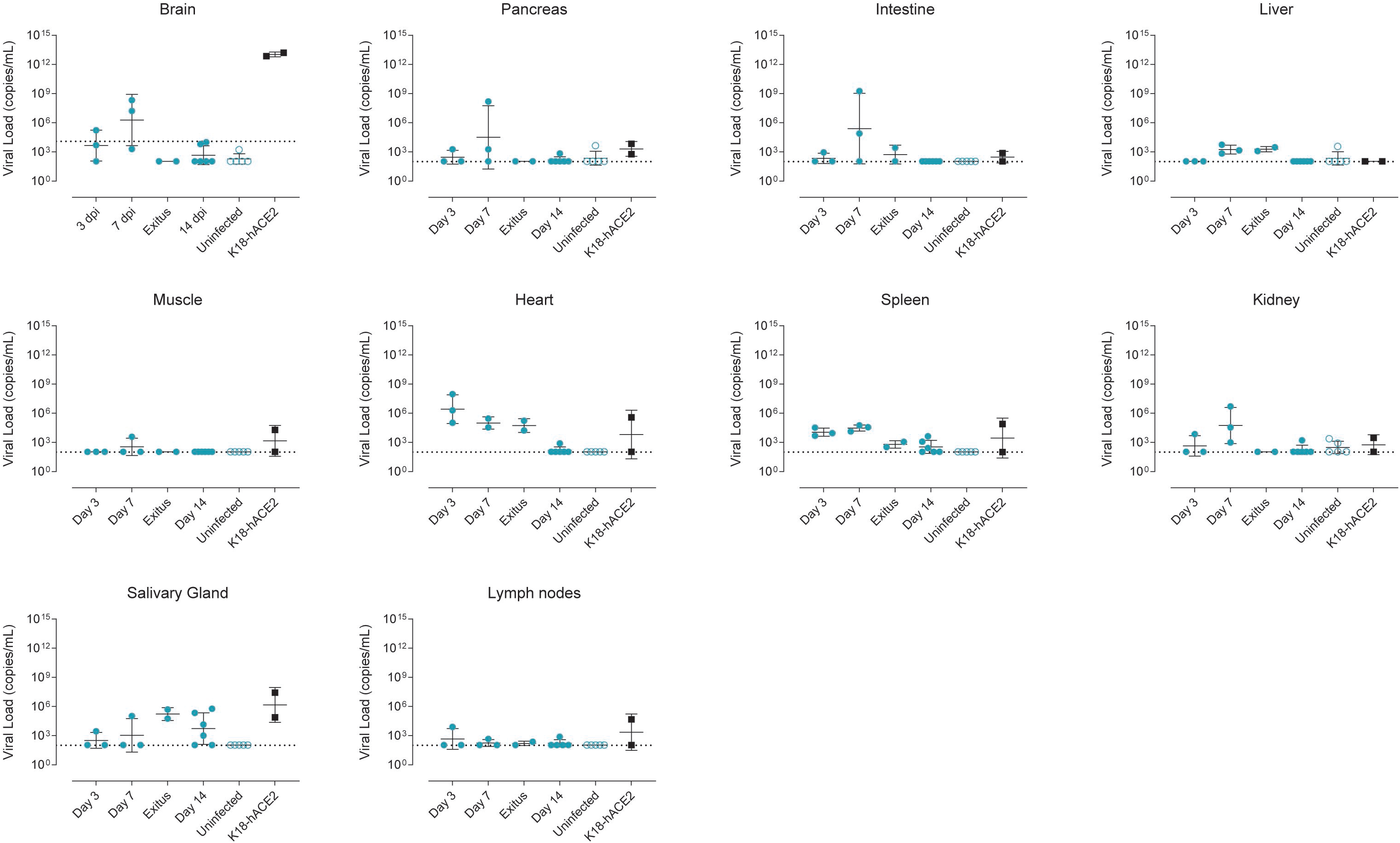
Tissue Tropism of SARS-CoV-2 in Col1a1-K18-hACE2 and K18-hACE2 Mice. SARS-CoV-2 viral RNA loads (copies/mL) were analyzed by RT-qPCR in Col1a1-K18-hACE2 mice tissues collected at 3,7 and 14 dpi (n=3 per timepoint) or upon fulfilment of humane endpoint criteria (n=1 at 8 dpi, n=1 at 9 dpi). K18-hACE2 tissues were collected at endpoint at 6 (n=1) and 7 dpi (n=1). Solid lines show the mean ±SD. Dashed line represents limit of detection, established by 2SD of uninfected animals.

### Col1a1-K18-hACE2 KI mice do not develop massive brain infection

To further investigate the level of viral infection in the brain (**Figure 5**), we performed a longitudinal virological and histological analysis of brain samples in all animals (n=32). Considering the full set of data, VL was detected at low levels in 3 of 8 Col1a1-K18-hACE2 mice at 3dpi and 2 out of 8 Col1a1-K18-hACE2 KI mice at 7 dpi. No VL was found in the additional animal that reached the humane endpoint. In contrast, all K18-hACE2 brain samples showed significantly higher VL at euthanasia (6-7dpi) (**Figure 6A**). However, no infective virus could be found in any Col1a1-K18-hACE2 mice at any timepoint by viral titration, while all K18-hACE2 presented high viral titers at 6-7dpi (**Figure 6B**).

**Figure 6.**
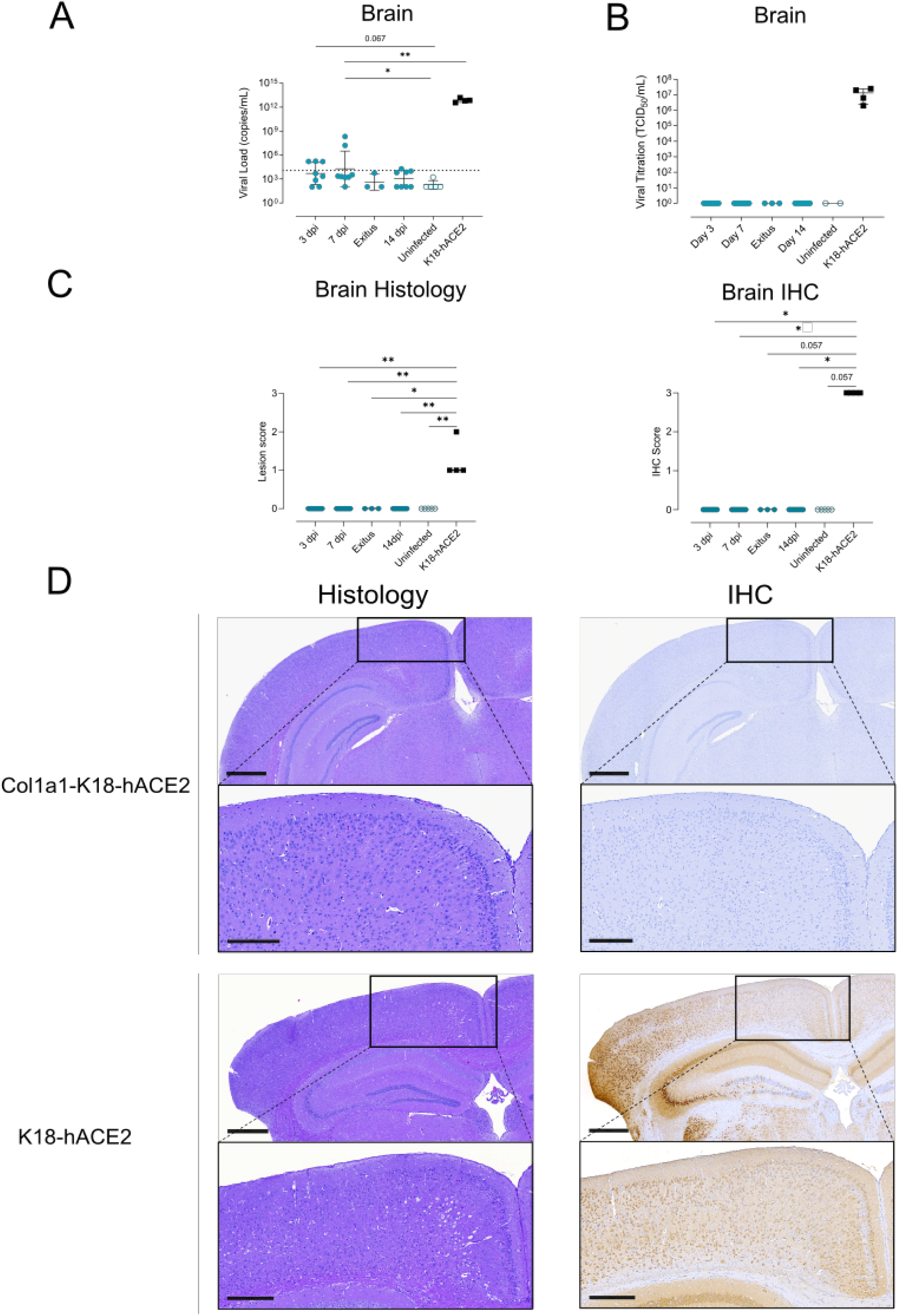
Progression of SARS-CoV-2 infection in brain from the Col1a1-K18-hACE2 KI mice. Col1a1-K18-hACE2 (blue circles, *n=27*) and K18-hACE2+ mice (black squares, *n=4*) were inoculated with 1000 TCID_50_ of a B.1 SARS-CoV-2 isolate (full shapes) or uninfected (empty shapes, *n=5*) and followed until 14 dpi. Samples were collected at 3, 7, 14 dpi and endpoint. (A) SARS-CoV-2 viral RNA loads (copies/mL) in brain extracts. Dashed line represents limit of detection, established by 2SD of uninfected animals. Statistical differences were identified using a Peto-Peto Left-censored samples test with correction for multiple comparisons. Solid line and bars represent mean and SD. (B) Viral titration of replicative virus (TCID_50/_mL) in brain samples from B.1 infected mice at different endpoints in Vero E6 cells on day 5 of culture. Solid line and bars represent mean and SD. (C) Brain histopathological scoring of multifocal lymphoplasmacytic meningo-encephalitis in brain (Left) and SARS-CoV-2 NP IHC scoring (right) in brain of both models. Statistical differences were identified using an Independence Asymptotic Generalized Pearson Chi-Squared Test for ordinal data (*p < 0.05; **p < 0.01). (D) Representative images of brain Histology and IHC images of both hACE2+ mouse models. Images show low-power magnification (top; bars: 600µm) and medium-power magnification (bottom; bars: 300μm). Col1a1-K18-hACE2 KI mice samples shown were collected at 7 dpi, while K18-hACE2 samples were collected at humane endpoint (6-7dpi).

For histological analysis, formalin-fixed brain samples were scored in a blinded fashion as described above. Histopathological analysis of the brain in the Col1a1-K18-hACE2 model showed no lesions at any of the analyzed timepoints nor those animals that required humane endpoint (**Supplementary** Figure 2). In contrast, brain samples from K18-hACE2 infected animals showed lesion scores between 1 and 2 **(Figure 6C,Left).** Furthermore, no detection of SARS-CoV-2 NP antigen in the brain of Col1a1-K18-hACE2 mice was observed by IHC at any timepoint **(Figure 6C)** or at the humane endpoint **(Supplementary** Figure 1); while K18-hACE2 mice showed a high density of antigen diffused throughout the tissue (score 3) at euthanasia **(Figure 6C, Right).** Brain changes in the transgenic model were consistent with the widely reported pathology induced by initial SARS-CoV-2 variants (12, 14, 21) and were observed alongside the detection of viral antigen in the brain (**Figure 6D**).

## Discussion

In this study, we characterized a novel hACE2 KI murine model by evaluating the course of infection with the B.1 D614G SARS-CoV-2 variant with a direct comparison to the well-described K18-hACE2 transgenic mouse model, in which this variant is known to be highly pathogenic. The most relevant findings included the susceptibility of the KI model to SARS-CoV-2 infection and the reduced lethality imputed to the reduced viral replication and lesions in the brain.

The development of this KI model was started at the beginning of the COVID-19 pandemic as an alternative to the K18-hACE2 transgenic mouse model. Even though the transgenic model had already been established and used in previous SARS-CoV studies (9), the availability of these animals was limited at first, as colonies needed to be restarted from cryopreserved embryos. In this context, several laboratories developed new hACE2 murine models for SARS-CoV-2 infection. Expression of the hACE2 transgene was achieved through various methods, including directed insertion using CRISPR/Cas9 technology (20) using different promoters such as Hepatocyte nuclear factor-3/forkhead homologue 4 (Hfh4) promoter (5) or the mouse ACE2 promoter (6), and by infecting mice with replication-defective adenovirus encoding hACE2 (3). In this work, we followed a different strategy wherein a single copy of the transgene was inserted into the COL1A1 locus, but under the K18 promoter as the original transgenic K18-hACE2 model. The selection of this locus was based on its known ability to provide reliable and ubiquitous expression of inserted sequences (19). Additionally, RMCE allowed the insertion of a single transgene copy which should better mimic physiological expression, potentially impacting lethality through reduction of the ectopic expression and modification of tissue distribution of the hACE2 receptor. A recent study identifying a correlation between lower hACE2 copy number and reduced pathogenicity further supports our initial hypothesis (22).

When compared with K18-hACE2 transgenic mice, B.1 infection of the new Col1a1-K18- hACE2 KI mouse model resulted in much less severe clinical signs with better responsiveness to stimuli, better physical appearance and no neurological signs. These observations were consistent with a lower viral replication in brain of Col1a1-K18-hACE2 KI mice, as summarized in **Supplementary** Figure 3. Focusing on the respiratory tract, viral replication was observed in both models, although VL and antigen detection were in general lower at 7 dpi in Col1a1-K18-hACE2 KI mice. In contrast, lung tissue lesions and inflammation were fully comparable in both models at this timepoint (**Supplementary** Figure 3).

To explain the different pathogenicity between each model, hACE2 expression was analyzed in different tissues. At mRNA level, expression in the lungs and nasal turbinates was comparable or higher in Col1a1-K18 hACE2 mice, while the brain and liver showed lower hACE2 expression, as reported for other transgenic approaches (22, 23). Expression of hACE2 protein showed additional differences; it confirmed lower expression in brain and liver and higher expression in pancreas of Col1A1-K18-hACE2 animals, but revealed higher expression in lung and spleen. In general, brain hACE2 expression in this new KI model seems to be closer to the physiological situation in humans, which showed low expression in the CNS, based on several techniques including transcriptomics and IHC (24). Among the clinical parameters analyzed in Col1a1-K18-hACE2 KI infected mice, we found that initial weight-loss was similar to that of the K18-hACE mice but was not accompanied by the increased severity of clinical signs that characterizes infection in these transgenic mice (neurological signs, lack of responsiveness, and poor general appearance) (13). Consistent with this observation, infection of Col1a1-K18-hACE2 KI mice led to partial recovery in weight by 14 dpi in most animals. Importantly, the survival rate of Col1a1-K18-hACE2 KI mice was over 70%, with three euthanized animals exclusively due to a weight loss of more than 20%, which was one of the humane endpoints in our study. These data are in line with other hACE2 KI mouse models in which substantial viral replication within the upper and lower respiratory tracts with limited spread to extrapulmonary organs has been described (8, 22, 25).

Among the tissues analyzed in Col1a1-K18-hACE2 KI infected mice, we found significant replication of SARS CoV-2 in the respiratory tract, mainly in the lungs. Despite viral clearance in Col1a1-K18-hACE2 KI mice which was confirmed by viral titration and IHC, lung lesions were similar between both models by 7dpi and remained after 14 dpi in KI animals recovering from the infection. The impact of viral replication on lung inflammation in Col1a1-K18-hACE2 was confirmed and found to be comparable to the inflammation profile reported in K18-hACE2 transgenic mice (21, 26). Infection of Col1a1-K18-hACE2 KI mice induced high levels of IFNγ, IL-6, IP-10, MCP-1 and MIP-1β cytokines. The longitudinal analysis showed early impact on IL-6 and IP-10 levels (3 dpi) and a clear trend to normalization by 14 dpi, with some residual inflammation remaining. This is consistent with the presence of lesions at this timepoint, and the immune cell infiltration described by 21 dpi in K18-hACE2 transgenic mice surviving infection (21).

The main difference between the Col1a1-K18-hACE2 KI and the K18-hACE2 transgenic models was found in neuroinvasion. Residual levels of viral RNA but no replication competent virus or viral proteins were detected in the brains of Col1a1-K18-hACE2 KI mice. Furthermore, no detectable brain lesions, such as the multifocal lymphoplasmacytic meningoencephalitis reported in K18-hACE2 transgenic mice (14) were detected by histology, and no SARS-CoV-2 nucleoprotein could be evidenced by IHC. Importantly, the three animals that reached humane endpoints did not present any evidence of brain viral replication but showed enhanced lung infection. Considering that the expression of hACE2 is lower in brain and higher in lung in Col1a1-K18-hACE2 compared to K18-hACE2, it seems that the targeted insertion of the hACE2 transgene can be a crucial factor leading to lower levels of brain infection and neurological clinical signs, as well as enhanced respiratory affection in this new model. However, different studies have shown that neuroinvasion is only partially dependent on the expression of hACE2 receptor (12), and that both delivery route and viral dose can play a role in the magnitude of the neuroinvasion (23, 27). In our case, to exclusively analyze the effect of receptor expression, we kept the intranasal administration route and TCID_50_ dose established and characterized in our own and others’ studies to ensure comparability of the results (13, 28).

Our study is a preliminary characterization of a new animal model and has consequently several limitations. The first limitation was the age of the animals used. The Col1a1-K18- hACE2 mice ranged in age from 5 to 11 months, depending on availability at the time of the study. However, although pathogenicity of SARS-CoV-2 infection has the potential to be higher in older animals, we could not detect significant age differences between euthanized and convalescent animals (mean+/-SD age of 9.1+/-2.0 and 8.0+/-2.1 months, respectively). Second, the expression of hACE2 has been analyzed at both the mRNA and protein levels in bulk tissues, but a further characterization in specific cell types will be necessary to link expression levels with pathology. Moreover, we have focused on the characterization of one viral variant and a single dose. The B.1 variant was selected for its demonstrated severity in K18-hACE2 mice and its inability to use mACE2 as an entry receptor, unlike B.1.351/Beta and Omicron BA1.1 variants (13, 15). While ancestral and earlier variants (B.1.1.7/Alpha, B.1.351/Beta B.1.617.2/Delta) were highly pathogenic in K18-hACE2 mice, Omicron BA.1.1 caused a milder infection with no weight loss nor neuroinvasion, therefore reducing lethality in this model (13). However, more recent Omicron subvariants such as BQ.1.1, BA.5 and XBB.1.5 appear to be regaining pathogenicity in K18-hACE2 mice (29–31). Increased lung infection, pro-inflammatory cytokines, and lung pathology are observed with these variants, with data on neuroinvasion and neurovirulence on the latter (31). Similarly, lower omicron variants has been described in KI mice (25, 32). Nevertheless, for the B.1 SARS-CoV-2 variant the Col1a1-K18-hACE2 model shows similarity with GSH in viral dynamics and pathology, with minimal neuroinvasion. This new model, however, has the upside of a wider reagent availability, scarce in GSH (1), and easier animal facility allowance.

In summary, several mouse models for the evaluation of antivirals and vaccines have been developed to date, with disease phenotypes ranging from mild to severe COVID-19-like condition. None of these models, however, can fully recapitulate all aspects of the disease as it occurs in humans. The development and further characterization of new animal models, like the Col1a1-K18-hACE2 model, may overcome some of these limitations and provide valuable tools to study certain aspects of COVID-19. Survival of Col1a1-K18-hACE2 KI mice after infection with a highly pathogenic variant could provide a model which better mimics human disease progression and thus could be instrumental for the study of specific disease aspects such as the consequences of inflammation (including loss of taste or smell) post-COVID conditions (PCC) or sequelae, long-term effects of drug therapies, and susceptibility to reinfection.

## Materials and methods

### Creation of the Knock-in Model

A new KI mouse model, **B6;129S-Col1a1^tm1(K18-hACE2)Irb^/Irsi** (Col1a1-K18-hACE2), was created by inserting the original K18-hACE2 transgene into the collagen COL1A1 locus using a recombinase-mediated cassette exchange (RMCE) FLP-FRT system in KH2 cells (19). The actual insertion site lies approximately 0.3 kb downstream of the 3’UTR end of COL1A1. As such, the inserted transgene remains identical to the original designed one (9) and the hACE2 cDNA is under the control of the K18 promoter rather than COL1A1. Specifically, the plasmid PGK-ATG-Frt was digested with EcoRV, and the sequence ATCAGACGTCGCTAGCGGCGCGCCGGTACTAGT was inserted to create a multi-cloning site.

The plasmid containing hACE2 under the control of the K18 promoter (K18-hACE2), supplied by Paul McCray, was digested with HpaI and XbaI enzymes. The resulting transgene fragment was isolated and purified, and then cloned into the EcoRV-NheI sites of the modified PGK-ATG-Frt plasmid to generate the targeting vector. The targeting vector was then co-transfected with an Flp expression construct into KH2 cells by electroporation. The cells were then placed under hygromycin selection for 9 days, after which drug resistant colonies were picked, expanded, and screened for the presence of the targeted transgene. Once confirmed, the modified embryonic stem cells were injected into mouse blastocysts. These injected mouse blastocysts were then placed into recipient foster females via embryo transfer techniques.

### Biosafety Approval and Virus Isolation

The execution of SARS-CoV-2 experiments was approved by the biologic biosafety committee of Germans Trias i Pujol Research Institute (IGTP) and performed at the Biosafety Level 3 laboratory (BSL-3) of the Comparative Medicine and Bioimage Centre (CSB-20-015- M8; CMCiB, Badalona, Spain).

The B.1 SARS-CoV-2 isolate used in this study was isolated from nasopharyngeal swabs of hospitalized patients in Spain as described elsewhere (33, 34). Briefly, viruses were propagated in Vero E6 cells (CRL-1586; ATCC, Virginia, VA, United States) for two passages and recovered by supernatant collection. The sequence of the SARS-CoV-2 variant tested is deposited at the GISAID Repository with accession IDs EPI_ISL_510689. EPI_ISL_510689 was the first SARS-CoV-2 virus isolated in Catalonia in March 2020 and, compared to the Wuhan/ Hu-1/2019 (WH1) strain, this isolate had the S protein mutations D614G, which is associated with the B.1 lineage, and R682L. Viral stocks were titrated on Vero E6 cells to use equivalent TCID50/mL using the Reed-Muench method and sequential 1/10 dilutions of the viral stocks as described previously (34).

### Animal Procedures and Study Design

All animal procedures were approved by the Committee on the Ethics of Animal Experimentation of the IGTP and were authorized by the *Generalitat de Catalunya* (code: 11222). All animal experiments followed the principles of animal welfare and the 3Rs. All experiments and sample processing were performed inside the BSL-3 facility. Col1a1-K18- hACE2 hemizygous KI mice were produced and bred at Parc Científic de Barcelona (PCB) by pairing hemizygous males for Tg(K18-ACE2)2Prlmn (or K18-hACE2) with non-carrier C57Bl6/J females. Animals used as control, B6.Cg-Tg(K18-ACE2)2Prlmn/J (or K18-hACE2) hemizygous transgenic mice (034860, Jackson Immunoresearch, West Grove, PA, United States) were bred at CMCiB in the Specific Pathogen Free Area (SPF) by pairing hemizygous K18-hACE2 males with non-carrier C57Bl6/J females. The genotype of the offspring regarding both K18-hACE2 and Col1a1-K18-hACE2 mice was determined by qPCR at the IGTP’s Genomics Platform from tail samples. Both animal models were kept in the BSL-3 facility during the whole experiment including acclimatization period. The housing conditions in the BSL-3 room were maintained as follows: a temperature of 22±2°C, humidity levels between 30-70%, 20 ACH, a 12h dark/light cycle, and access to food and water *ad libitum*.

A total of 36 adult mice aged 5-11 months were used in this experiment, consisting of 32 KI Col1a1-K18-hACE2 mice and 4 transgenic K18-hACE2 mice. All groups were sex balanced. The infections were performed in two separate experiments between January and June 2022. Mice were anaesthetized with isoflurane (FDG9623; Baxter, Deerfield, IL, USA) and infected with a B.1 isolate (27 Col1a1-K18-hACE2 and 4 K18-hACE2 mice). The uninfected control group received PBS (5 Col1a1-K18-hACE2).

Infection was performed using 1,000 TCID_50_ of B.1 SARS-CoV-2 isolate in 50 μl of PBS (25 μL/nostril), or PBS only (25 μL/nostril) for the control group. All mice fully recovered from the challenge and anaesthesia procedures. Following the challenge, body weight and clinical signs were monitored daily. Eight animals per group were euthanized at days 3, 7, 14 dpi or upon fulfilment of human endpoint, for viral RNA quantification and histological analyses (**Supplementary Table 2**). The endpoint criteria for animal welfare were established based on a body weight loss of more than to 20% of the initial body weight and/or the display of moderate to severe clinical signs (including neurological signs), in accordance with previous studies (12–14). The evaluated clinical signs included respiration, physical appearance, lack of responsiveness and neurological signs, that were scored from 0-2 depending on severity (**Supplementary Table 1**). Euthanasia was performed under deep isoflurane anaesthesia by whole blood extraction via cardiac puncture and cervical dislocation. Oropharyngeal swab, lung, brain, and nasal turbinate were collected for viral and hACE2 RNA quantification, histological and IHC analyses. For the latter techniques, tissues were fixed by immersion in 10% buffered formalin. An additional set of 9 tissues was collected to further characterize the KI model both for viral RNA quantification and hACE2 receptor expression. These tissues included muscle, intestine, liver, kidney, lymph nodes, spleen, heart, pancreas, and salivary glands.

### Tissue sampling and processing

The collected tissues were processed as described by Tarrés-Freixas et al. 2022 (13). Briefly, approximately 100 mg of each tissue was collected in 1.5 mL Sarstedt tubes (72607; Sarstedt, Nümbrecth, Germany) containing 500 μL of DMEM medium (11995065; ThermoFisher Scientific) supplemented with 1% penicillin–streptomycin (10378016; ThermoFisher Scientific). A 1.5 mm Tungsten bead (69997; QIAGEN, Hilden, Germany) was added to each tube, and samples were homogenized twice at 25 Hz for 30 s using a TissueLyser II (85300; QIAGEN) before being centrifuged for 2 min at 2,000 × g. Supernatants were then stored at −80°C until the analysis of hACE2 expression and VL by RT-qPCR.

### hACE2 mRNA Expression

RNA tissue extraction was performed by using the Viral RNA/ Pathogen Nucleic Acid Isolation kit (A42352; ThermoFisher Scientific), optimized for use with a KingFisher instrument (5400610; ThermoFisher Scientific), following the manufacturer’s instructions. The expression of hACE2 receptor in tissue was detected by RT-qPCR with a predefined TaqMan assay with a ACE2 target (Hs01085333_m1, ThermoFisher Scientific). Mouse *gapdh* gene expression was measured in duplicate for each sample using TaqMan®gene expression assay (Mm99999915_g1; ThermoFisher Scientific) as amplification control. Data was graphed both as the delta-Ct values of hACE2 minus GAPDH for each tissue, and the relative expression of the receptor (delta-delta Ct value) of Col1a1-K18-hACE2 minus K18-hACE2 tissues.

### ACE2 Protein Expression

ACE2 protein expression was assessed by WB. Tissue samples were obtained from uninfected Col1a1-K18-hACE2, K18-hACE2 and WT mice and flash frozen. Frozen sections of each tissue were sliced by cryostat processing (Leica CM1950). and homogenized in 100uL of RIPA buffer for 15 min on ice, followed by centrifugation at 1000g for 10 min. Supernatant was then isolated, and total protein concentration was quantified by Pierce™ BCA Protein Assay (23225; ThermoFisher Scientific) following manufacturer’s instructions.

For WB, 3 µg of lysate total protein (as quantified Pierce™ BCA Protein Assay (23225; ThermoFisher Scientific) were run in NuPAGE Bis-Tris 4-12% acrylamide gels (NP0321; ThermoFisher Scientific). BlueStar pre-stained protein ladder was used as a molecular weight marker (523005; NIPPON Genetics, Tokyo, Japan). Proteins were blotted onto PVDF membrane (Ref. 1704156; Bio-Rad) using the Trans-Blot Turbo Transfer System (Ref. 1704150 Bio-Rad) and membranes were blocked with EveryBlot Blocking Buffer (12010020; Bio-Rad). Primary antibodies used were: Anti-hACE2 specific monoclonal (1:1000; AB108209, Abcam) (20), Anti-ACE2 polyclonal (1:1000 AF933; R&D Systems) and Anti-GAPDH (1:5000; Ab9485; Abcam). Secondary antibodies were: donkey anti-Rabbit (1:10000; Ref. 711-036-152; Jackson Immunoresearch) or a donkey anti-Goat (1:10000; Ref. 705-035- 147; Jackson Immunoresearch). Membranes were developed with SuperSignal™ West Femto Maximum Sensitivity Substrate (Ref. 34094; ThermoFisher Scientific) according to the manufacturer’s instructions and acquired with ChemiDOC^TM^ MP Imaging System (12003154; Bio-Rad). Images were processed and merged with ImageLab 6.0.1 software (Bio-Rad).

### SARS-CoV-2 PCR Detection and Viral Load Quantification

Viral RNA was quantified by RT-qPCR in both the standard set of samples (oropharyngeal swab, lung, brain, and nasal turbinate), and the extended set (muscle, intestine, liver, kidney, pancreas, salivary glands, lymph nodes and spleen).

RNA extraction was performed by using the Viral RNA/ Pathogen Nucleic Acid Isolation kit (A42352; ThermoFisher Scientific), optimized for use with a KingFisher instrument (5400610; ThermoFisher Scientific), following the manufacturer’s instructions. PCR amplification was based on the 2019-Novel Coronavirus Real-Time RT-PCR Diagnostic Panel guidelines and protocol developed by the American Center for Disease Control and Prevention (CDC-006-00019, v.07). Briefly, a 20 μL PCR reaction was set up containing 5 μL of RNA, 1.5 μL of N2 primers and probe (2019-nCov CDC EUA Kit; catalog number 10006770; Integrated DNA Technologies, Coralville, IA, USA) and 10 μl of GoTaq 1-Step RT-qPCR (Promega, Madison, WI, USA). Thermal cycling was performed at 50°C for 15 min for reverse transcription, followed by 95°C for 2 min and then 45 cycles of 95°C for 10 s, 56°C for 15 s and 72°C for 30 s in the Applied Biosystems 7,500 or QuantStudio5 Real-Time PCR instruments (ThermoFisher Scientific). For absolute quantification, a standard curve was built using 1/5 serial dilutions of a SARS-CoV-2 plasmid (2019-nCoV_N_Positive Control; catalog number 10006625, 200 copies/μL, Integrated DNA Technologies), which was run in parallel with all PCR determinations. Viral RNA from each sample was quantified in triplicate, and the mean viral RNA concentration (in copies/mL) was extrapolated from the standard curve and corrected by the corresponding dilution factor. Mouse *gapdh* gene expression was measured in duplicate for each sample using TaqMan®gene expression assay (Mm99999915_g1; ThermoFisher Scientific) as amplification control.

### Viral Titration

Lung tissues were evaluated for the presence of replicative virus by titration in Vero E6 cells as previously described (14, 34, 35). Briefly, after tissue homogenization, each sample underwent sequential 10-fold dilutions in duplicate, transferred onto a monolayer of Vero E6 cell in a 96-well plate, and incubated at 37°C and 5% CO_2_. Plates were monitored daily under a microscope, and at 5 dpi, wells were evaluated for the presence of cytopathic effects. The amount of infectious virus was calculated by determining the TCID50 using the Reed–Muench method.

### Histological and Immunohistochemical Analyses

Tissue samples were recovered at the designated endpoint (3,7,14 dpi or humane endpoint) and fixed by immersion in 10% buffered formalin. Lung, nasal turbinate, and brain samples were routinely processed for histological examination, with haematoxylin & eosin-stained slides examined under an optical microscope in a blinded fashion. Brain was cut into two coronal sections, including cerebellum/pons and hemispheres at thalamus level. A semi-quantitative approach based on the amount of inflammation (none, mild, moderate, or severe) was used to assess the damage caused by SARS-CoV-2 infection in mice, following a previously published scoring system (14, 36). Additionally, an IHC technique was employed to detect SARS-CoV-2 NP antigen in nasal turbinate, lung, and brain sections from all animals, using a rabbit monoclonal antibody (40143-R019, Sino Biological, Beijing, China) at a 1:15,000 dilution. The amount of viral antigen in tissues was semi-quantitatively scored in a blinded fashion (low, moderate, and high amount, or lack of antigen detection) (14, 36).

### Cytokine Quantification

To assess the viral-driven inflammation in lung in both animal models, the levels of IP-10, IL-6, IFNγ, MCP-1 and MIP-1β cytokines were analyzed by Luminex in tissue extracts. Lung samples were processed as described above and stored at -80°C until analysis. In Col1a1- K18-hACE2 mice, cytokines were analyzed at 3,7,14 dpi and those euthanized by humane endpoint criteria (exitus). Uninfected Col1a1-K18-hACE2 and K18-hACE2 infected animals were used as reference groups.

Cytokines were measured by Luminex xMAP technology and analyzed with xPONENT 3.1 software (Luminex Corporation) using the MCYTOMAG-70 kit, according to the manufacturers’ protocol with minor modifications. Briefly, after cytokine staining, samples underwent an overnight incubation on a rocking shaker at 4°C, using 2% PFA to ensure complete inactivation of any remaining SARS-CoV-2 particles; a fixation that does not alter cytokine quantification (37). Before plate acquisition, the PFA was washed away and replaced with sheath fluid.

### Pseudovirus generation and neutralization assay

To assess the generation of neutralizing antibodies upon SARS-CoV-2 B.1 infection, we used a previously described pseudovirus-based neutralization assay (38). Briefly, in Nunc 96-well cell culture plates (Thermo Fisher Scientific), 200 TCID50 of a luciferase-reporter HIV-based pseudovirus bearing the BA.1 SARS-CoV-2 spike were preincubated with four-fold serial dilutions (1/60 - 1/61,440) of the heat-inactivated (56°C for 30 minutes) serum samples, for 1 hour at 37°C. Then 1x10^4^ HEK293T/hACE2 cells treated with DEAE-Dextran (Sigma-Aldrich) were added. Results were read after 48 hours using the EnSight Multimode Plate Reader and BriteLite Plus Luciferase reagent (PerkinElmer, USA). The values were normalized, and the ID50 (reciprocal dilution inhibiting 50% of the infection) were calculated as described (39).

### Statistical Analyses

All figures were generated using GraphPad Prism 9.0.0. Statistical analyses were performed using R v4.3. Survival Rates were estimated with Kaplan-Meier curves and compared with the Log-rank test. Datasets with an abundance of data below the limit of detection, like VL, were analyzed using the Peto-Peto Left-censored samples test with correction for multiple comparisons. Histopathological, IHC scores and viral titrations were compared using an Independence Asymptotic Generalized Pearson Chi-Squared Test for ordinal data. Cytokine titers were compared by a Kruskal Wallis, with pairwise comparisons conducted using Conover’s non-parametric test.

## Supporting information

Suppemental Figures

## Acknowledgements

The authors would like to acknowledge Jorge Díaz, Yaiza Rosales-Salgado, Rosa María Ampudia-Carrasco, Sergi Sunyé-Casas and Mireia Martínez from the CMCiB for their essential help in the BSL3 facility and the K18-hACE2 mouse colony. We also thank Marisa Larramona from Parc Científic de Barcelona for her invaluable help with the knock-in mouse colony. We thank the Dormeur Fondation for their financial support for the acquisition of the QuantStudio-5 real-time PCR system.

## Author Contributions

AP-G, FT-F, BT, and JB conceived and designed the experiments. AP-G, FT-F, and BT performed the animal procedures. AP-G, FT-F, MP, ER-M, DR-R, DP-Z, JM-B, EP, JS, BT performed the analytical experiments. AP-G, B, JS, JV-A, VU and JB analyzed and interpreted the data. SF, BTondelli established and provided the Knock-in mouse colony. SC established communication between animal facilities and contributed to the veterinary report verification to allow the knock-in animal shipment to the CMCiB facility. DP-Z, JM-B, DR-R, NI-U provided key reagents. AP-G, BT, and JB wrote the paper. All authors contributed to the article and approved the submitted version.

## Data availability

The data supporting the findings of this study are documented within the paper and are available from the corresponding authors upon request.

