## Supplementary material for "A human-ACE2 knock-in mouse model for SARS-CoV-2 infection recapitulates respiratory disorders but avoids neurological disease associated with the transgenic K18-hACE2 model": Suppemental Figures

**Running title**: B.1 SARS-CoV-2 infection of a new hACE-2 Knock-in mice

*Anna Pons-Grífols[1], Ferran Tarrés-Freixas[1,2,3,4], Mònica Pérez[2,3], Eva Riveira-Muñoz[1], Dàlia Raïch-Regué[1], Daniel Pérez-Zsolt[1], Jordana Muñoz-Basagoiti[1], Barbara Tondelli[5], Edwards Pradenas[1], Nuria Izquierdo-Useros[1,6,7,], Sara Capdevila[8], Júlia Vergara-Alert[2,3], Victor Urrea[1], Jorge Carrillo[1,6,7], Ester Ballana[1,6], Stephen Forrow[5], Bonaventura Clotet[1,4], Joaquim Segalés[2,9], Benjamin Trinité[1,#], Julià Blanco[1,4,6,7,#].*

[1]IrsiCaixa, Can Ruti Campus, Badalona, Spain
[2]Unitat mixta d'investigació IRTA-UAB en Sanitat Animal, Centre de Recerca en Sanitat Animal (CReSA), Campus de la Universitat Autònoma de Barcelona (UAB), Bellaterra, Spain
[3]IRTA, Programa de Sanitat Animal, Centre de Recerca en Sanitat Animal (CReSA), Campus de la Universitat Autònoma de Barcelona (UAB), Bellaterra, Spain
[4]University of Vic-Central University of Catalonia (UVic-UCC), Vic, Spain
[5]Mouse Mutant Core Facility (PBA2B2), Institute for Research in Biomedicine (IRB), Parc Científic de Barcelona, Barcelona, Spain
[6]Germans Trias i Pujol Research Institute (IGTP), Badalona, Spain
[7]CIBER Infectious Diseases (CIBERINFEC), Carlos III Health Institute, Madrid, Spain.
[8]Comparative Medicine and Bioimage Centre of Catalonia (CMCiB), Germans Trias i Pujol Research Institute (IGTP), Badalona, Spain.
[9]Departament de Sanitat i Anatomia Animals, Facultat de Veterinària, Universitat Autònoma de Barcelona (UAB), Campus de la UAB, Bellaterra, Spain.

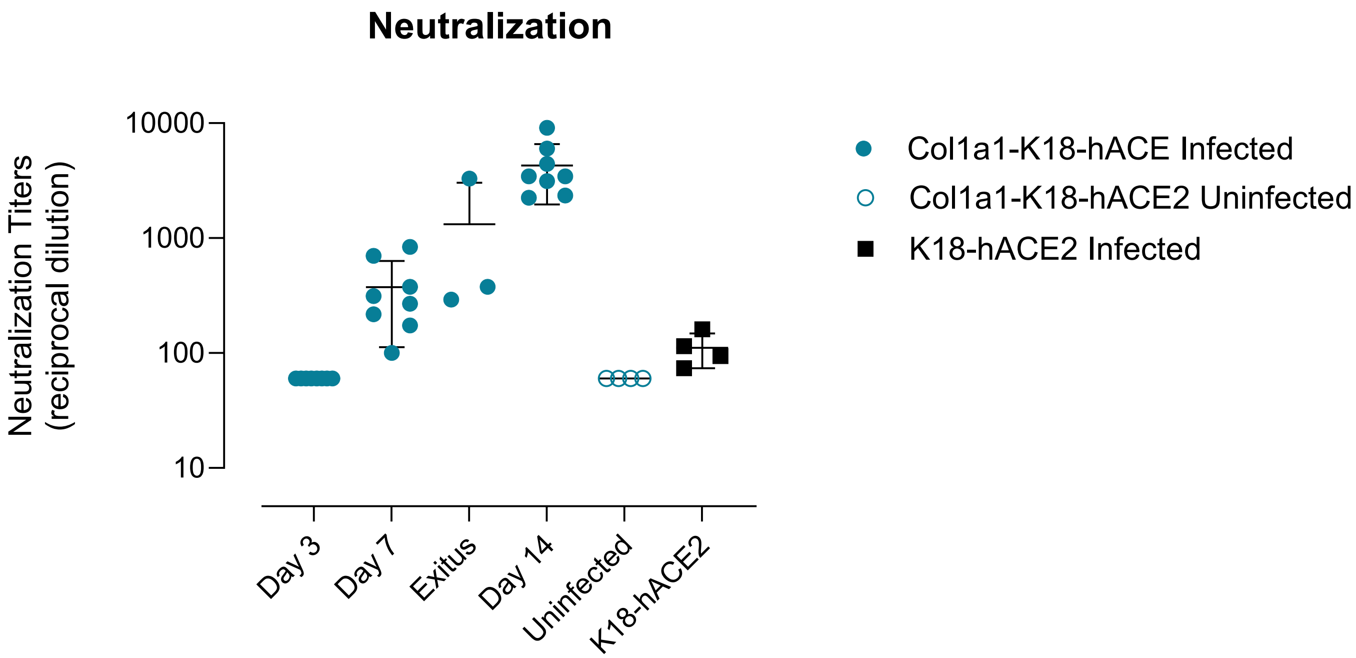


**Supplementary Fig 1. SARS-CoV-2 neutralization activity.** Serum samples from the indicated animals were assayed for their ability to neutralize a B.1 pseudovirus (see methods). Figure shows reciprocal dilutions of serum blocking 50% of pseudovirus infectivity for each animal. Geometric mean and confidence intervals are shown for each group.

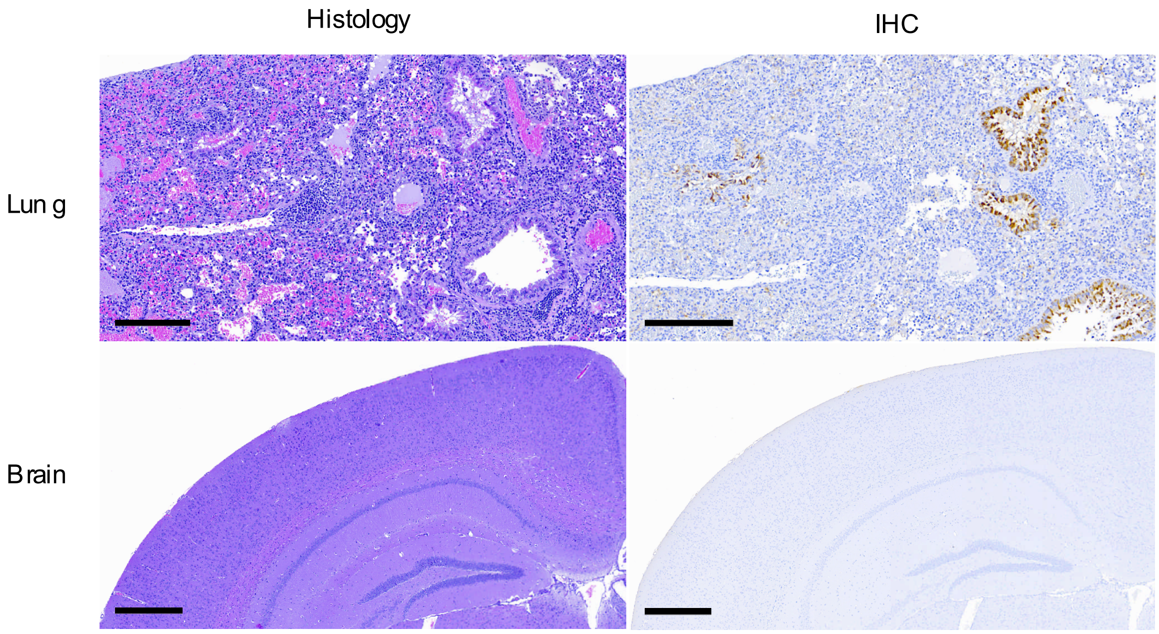


**Supplementary Figure 2. Pathological and Immunohistochemistry analysis of SARS-CoV-2 antigen in lung and brain of Col1a1-K18-hACE2 animals reaching humane endpoint criteria**. (Upper) Lung from mouse 9980-21 (euthanized at 9 dpi, 9.8 months old) displaying an area of severe bronchointerstitial pneumonia (left) and mild to moderate SARS-CoV-2 antigen mainly located in bronchiolar epithelium from different bronchiole (right). This animal had an overall pathological and immunohistochemical scores of 2. Images show medium-power magnification bars (200 µm). (Lower) Brain from mouse 012802-22 (euthanized at 10 dpi exitus, 6.27 months old) displaying normal histological features (left) and lack of SARS-CoV-2 antigen (right). This animal had overall pathological and immunohistochemical scores of 0. Images show low-power magnification bars (500 µm).

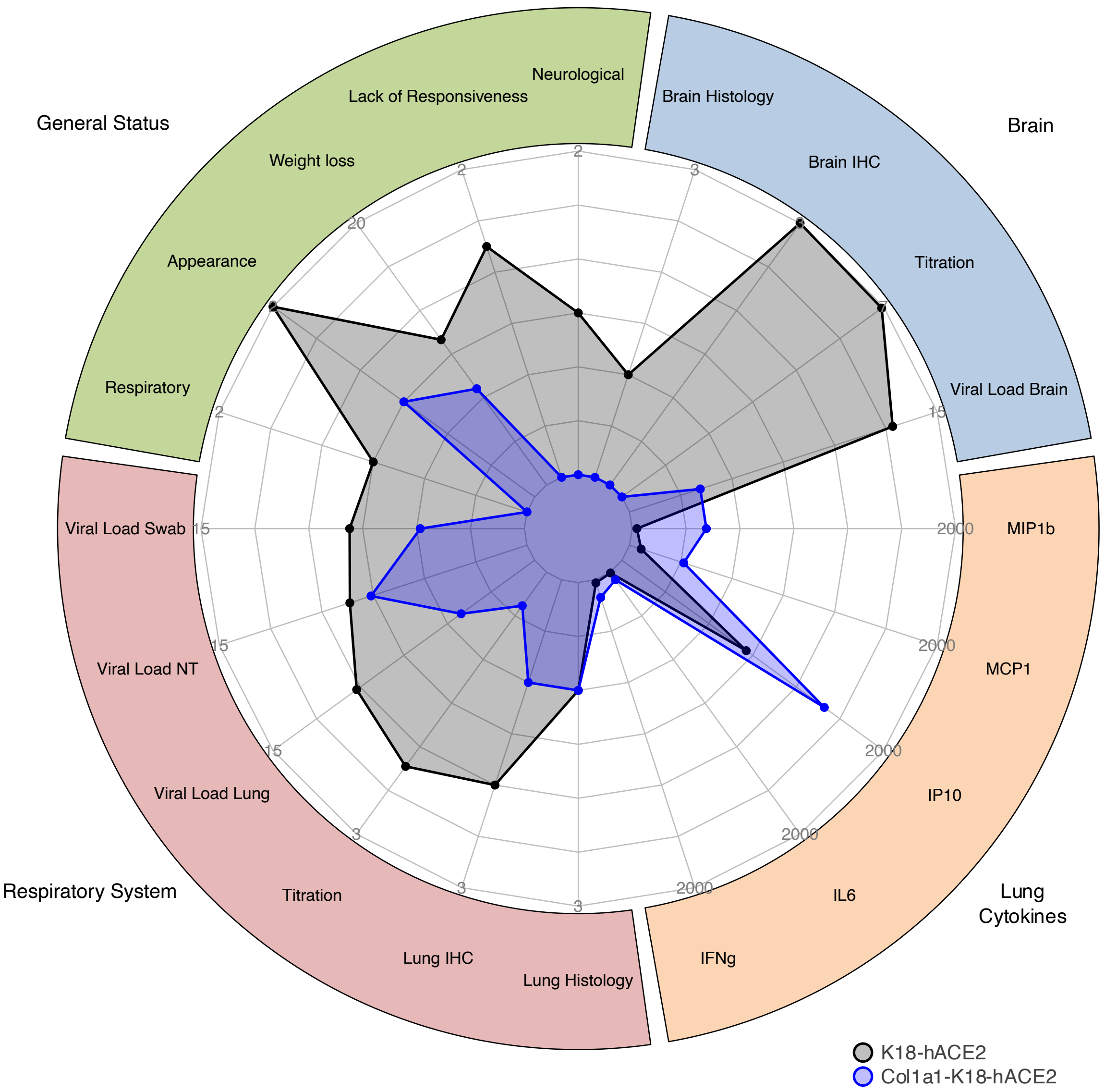


**Supplementary Fig 3. Spider Plot of clinical data compilation at 7 dpi from both animal models.** Comparative data summary of Col1a1-K18-hACE2 and K18-hACE2 mice infection by a B.1 SARS-CoV-2 isolate. Data at 7dpi of the KI (blue; euthanized at 7 dpi) and K18-hACE2 (black; euthanized at 6-7dpi) of different parameters was compiled and median values are shown in each axis of the spider plot. Data ranges are adjusted for each kind of data, and shown by concentric circles, being the outside circle the highest level. Clinical status was assessed in different categories:

respiration, appearance, weight-loss, lack of responsiveness and neurological signs. Histology and IHQ data were semiquantitatively scored from 0-2, 0-3 and 0-3 respectively as described in Materials and Methods section. Viral loads are shown as Log10 copies/mL, titration as Log10 TCID50 and cytokines as pg/mL.

A

| Clinical Status Score | 0 | 1 | 2 |
|---|---|---|---|
| Body weight-loss | <10% | 10-20% | >20% |
| Appearance | Normal | Some piloerection, possible hunched back, eyes not fully open, possible secretions | Majority Piloerected, hunched back, eyes half closed, possible secretions |
| Respiratory | Normal | Moderately reduced | Severely reduced, laboured |
| Lack of Responsiveness | Normal,responds to touch | Reduced mobility, moderate response to touch or auditory stimuli | No mobility/ little or no response to touch or auditory stimuli |
| Neurological | No signs | Impaired activity, possible tremors | Suppressed activity,tremors |

B

| | Col1a1-K18-hACE2 (KI) | | | | | | | | K18-hACE2 | | | | Median | |
|---|---|---|---|---|---|---|---|---|---|---|---|---|---|---|
| Clinical Status | IGTP-012801-22 | IGTP-012799-22 | IGTP-012800-22 | IGTP-012815-22 | IGTP-012816-22 | IGTP-009993-21 | IGTP-009992-21 | IGTP-009983-21 | IGTP-006365-21 | IGTP-006367-21 | IGTP-010256-21 | IGTP-010255-21 | KI | K18 |
| Body weight-loss | 1 | 1 | 1 | 1 | 1 | 0 | 0 | 0 | 1 | 1 | 1 | 1 | **1** | **1** |
| Appearance | 1 | 1 | 2 | 2 | 1 | 2 | 1 | 1 | 2 | 2 | 1 | 2 | **1** | **2** |
| Respiratory | 0 | 0 | 1 | 1 | 0 | 0 | 0 | 0 | 1 | 1 | 1 | 1 | **0** | **1** |
| Lack of Responsiveness | 1 | 1 | 0 | 0 | 0 | 0 | 0 | 0 | 1 | 1 | 2 | 2 | **0** | **1,5** |
| Neurological | 0 | 0 | 0 | 0 | 0 | 0 | 0 | 0 | 1 | 1 | 1 | 1 | **0** | **1** |

**Supplementary Table 1.** Clinical scoring system for monitoring of Col1a1-K18-hACE2 and K18-hACE2 models. (A) All categories were scored from 0-2, as pictured. (B) Clinical status data at 7dpi. Each column represents an individual animal. Col1a1-K18-hACE2 (n=8), K18-hACE2 (n=4, includes animals euthanized at 6dpi).

| | Days post- SARS-CoV-2 inoculation (dpi) | | | | | | |
|---|---|---|---|---|---|---|---|
| **Group** | **3** | 6 | **7** | 8 | 9 | 10 | **14** |
| Col1a1-K18-hACE2 *(n=27)* | 8 | - | 8 | 1* | 1* | 1* | 8 |
| K18-hACE2 *(n=4)* | - | 2* | 2* | - | - | - | - |
| Col1a1-K18-hACE2 Uninfected *(n=5)* | - | - | - | - | - | - | 5 |

**Supplementary Table 2**. Number of animals euthanized after SARS-CoV-2 inoculation. The study's established endpoints of 3,7 and 14 dpi are indicated in grey columns, with 8 animals sampled per timepoint. Euthanasias done under humane endpoint criteria are marked with an asterisk.
